# The mammary glands of cows abundantly display receptors for circulating avian H5 viruses

**DOI:** 10.1101/2024.05.24.595667

**Authors:** María Ríos Carrasco, Andrea Gröne, Judith M.A. van den Brand, Robert P. de Vries

## Abstract

Influenza A viruses (IAV) from the H5N1 2.3.4.4b clade are circulating in dairy farms in the United States of America, and goat infections have also been reported. These ruminants were presumed not to be hosts for IAVs. Previously, IAV-positive mammalian species were hunters and scavengers, possibly getting infected while feeding on infected birds. It is now presumed that H5N1 viruses that circulate in US dairy cattle transmit through a mammary gland route, in contrast to transmission by aerosols via the respiratory tract, although the latter cannot be excluded. The receptor display in the mammary and respiratory tract in cows for IAVs is unknown. Here, we used recombinant HA proteins representing current circulating and classical H5 viruses to determine the distribution of IAV receptors in the respiratory and mammary tract tissues of cows and goats. Most of the sialome of the cow and goat respiratory tract is lined with sialic acid modifications such as N-glycolyl and O-acetyl, which are not bound by IAV. Interestingly, the H5 protein representing the cow isolates bound significantly in the mammary gland, whereas classical H5 proteins failed to do so. Furthermore, whereas the 9-O-acetyl modification is prominent in all tissues tested, the 5-N-glycolyl modification is not, resulting in the display of receptors for avian IAV hemagglutinins. This could explain the high levels of virus found in these tissues and milk, adding supporting data to this possible virus transmission route.

## Introduction

Since late 2020, we have been experiencing an unprecedented global outbreak of high pathogenic H5Nx influenza A viruses (IAV) (1, 2). These viruses circulate year-round from the Northern to Southern hemisphere and cause incredibly high mortality in avian species with significant transmission to mammals (3-5). The transmission route to mammals is under debate as most infected mammals are hunters and scavengers, possibly getting infected while consuming bird remains (6-9). Seal infections, on the other hand, have been known to occur for decades due to their proximity to wild waterfowl (10); however, the spread into pinnipeds of the southern hemisphere was unprecedented. The appearance of the viruses in the Antarctic and the current zoonotic event to ruminants add even more novelties to this remarkable outbreak (11). The latest isolation of IAVs in cows and goats was an unexpected transmission event as these species were believed not to be hosts of IAVs, while they are hosts for influenza D viruses (IDV) (12, 13). However, it quickly became apparent that mammary tract tissues are probably essential in the transmission route, as milk samples most frequently contain high viral titers (14).

The molecular determinant of the current widespread H5Nx viruses is currently unknown. While the E627K mutation in the polymerase is commonly found in viruses isolated from mammalian species, the moment of occurrence is whether viruses with this mutation circulate in birds or is this mutation is immediately selected in mammals. Only the human isolate associated with the current cow outbreak contains this mutation (15). The HA gene has been remarkably stable with no mutations in the receptor binding site (RBS) in the 2.3.4.4b virus for years. Genotypes such as 2.3.4.4a,c, and e harbor some RBS mutations; however, these clades are minority species. Thus, it is likely that 2.3.4.4b virus HA proteins already have an optimal receptor binding specificity, allowing their panzootic nature.

Most IAVs use sialic acids (SIA) to enter a host cell. The glycome that presents these SIAs in the respiratory tract of farm animals such as goats and cows is poorly defined (16), in contrast to several other bovine proteins such as submaxillary gland mucins and fetuin (17-20). It is, however, known that ruminants display a set of modified SIAs that other IAV host species do not. These include the N-glycolyl modification at the C5 position (5-N-glycolyl), which is created by the cytidine monophospho-N-acetylneuraminic acid hydroxylase (CMAH) (21). CMAH is a mammalian-specific enzyme that is non-functional in various influenza hosts, including humans, ferrets, dogs, and seals (22). Cows and goats express a functional enzyme and thus abundantly display Neu5Gc (23) which is not a receptor for most influenza A viruses (24). The other abundant SIA modifications in the cow respiratory tract are O-acetyls, which can be present on the 4, 7, 8, and 9 positions. The latter is the essential receptor for influenza C, D (25), and a variety of coronaviruses (26, 27), but a non-ligand for influenza A virus (28-30). The current dogma is that these SIA modifications, not present in the avian reservoir, are decoy or blocking moieties for IAVs.

To determine if 2.3.4.4b H5 influenza A viruses can bind to available receptors in the respiratory and mammary tract of cows, we used formalin-fixed and paraffin-embedded (FFPE) nasal, tracheal, lung, and mammary gland tissues of cows and goats (31). The animals were submitted to the Veterinary Pathology Diagnostic Centre (VPDC) for postmortem evaluation. No animals were euthanized for this study.

## Results

### 2.3.4.4b viruses isolated from goats and dairy cows have a conserved receptor binding site

The receptor binding site, the HA of IAVs, is composed of the 190 helices (AA180-200), the 130-(AA130-140), and 220-(AA220-230) loops. Since the introduction of the 2.3.4.4b clade a decade ago, the receptor binding domain has remained relatively conserved (Fig 1). Significant changes in 2.3.4.4 viruses, compared to the classical A/Vietnam/1203/2004 (H5VN) and A/Indonesia/05/2005 (H5IN), include the 130 loop, the loss of a glycosylation site at position 158, and significant perturbations in the 190 helix and the 222 and 227 positions in the 220-loop that are directly involved in receptor binding (32, 33). Taking A/duck/France/161108h/2016 (H5FR) as a reference, we only observe minimal amino acid changes in the recent North American mammalian-derived viruses. The mutations observed are outside the canonical RBS, namely L122Q, T144A, T199I, and V214A (H3 numbering). Of note, some cow sequences isolated later now contain A160T and thus restore the glycosylation site at position 158. Conclusively, although the 2.3.4.4b H5 viruses have been circulating for several years around the globe in different hosts, the receptor binding domain remained conserved.

**Figure 1.**
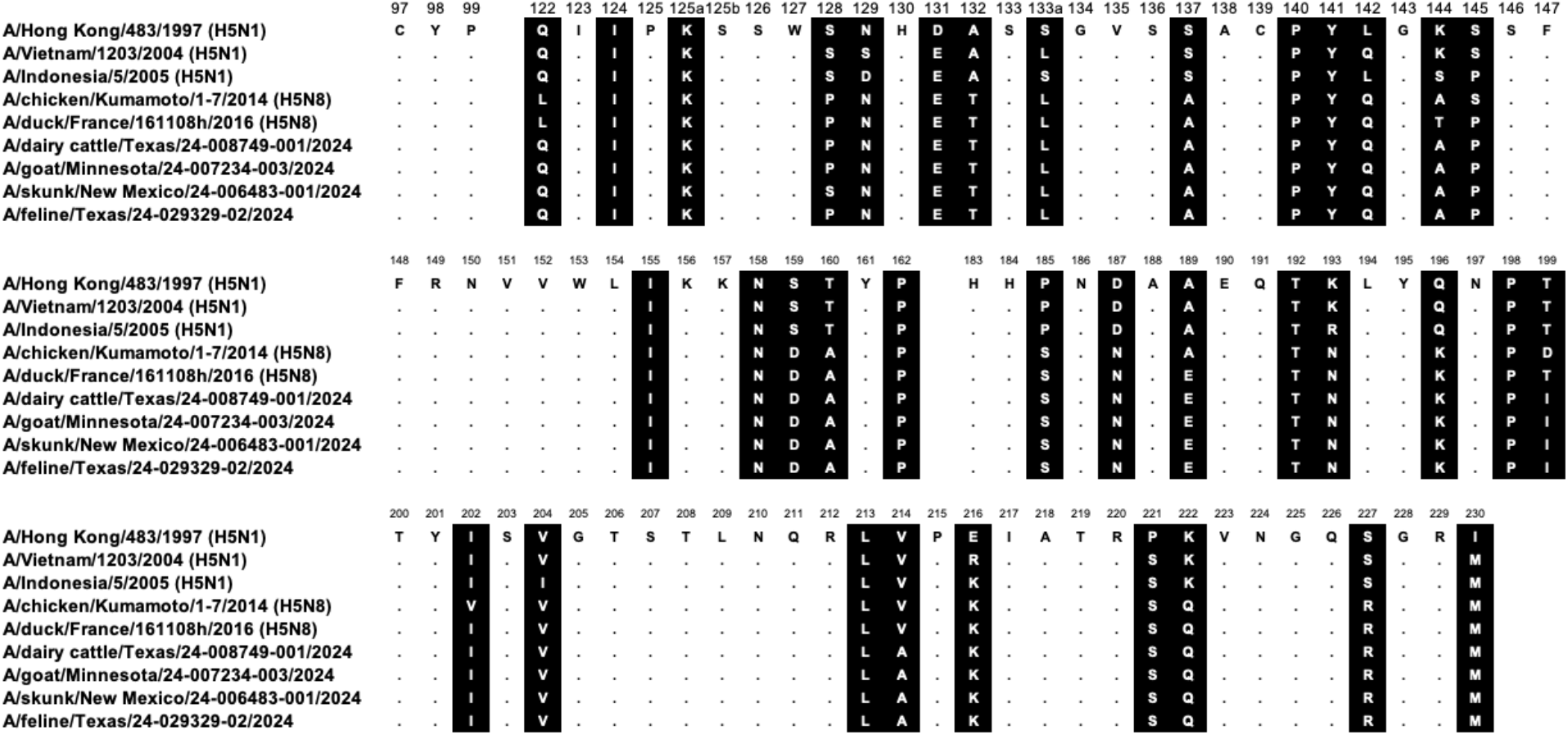
HA receptor binding site amino acid alignment of the H5 hemagglutinins used in this study. Alignment of the receptor binding site residues with amino acid positions (H3 numbering) indicated above the alignment, non-conserved residues highlighted in black, dots indicating identical amino acids. Several mammalian sequences isolate in North America are shown to demonstrate the close relationship.

### Contemporary 2.3.4.4b HA proteins bind efficiently to α2,3 linked Neu5Ac containing sialosides in mammary tract tissues of cows

A central observation of IAV-infected dairy cows is mammary gland infections (14). To determine if this gland has receptors for H5 IAVs to support active replication, we used both classical and contemporary H5 proteins for tissue binding studies. We used the classical H5 derived from A/Indonesia/05/2005 (H5IN), A/Vietnam/1203/2004 (H5VN), and the Y161A (H5VN Y161A) mutant that confers binding to Neu5Gc (34). To detect 9-O-acetylated structures, we employed an enzymatically inactive influenza D virus hemagglutinin esterase fusion protein (D/OK) (12). We also used commonly employed plant lectins SNA and MAL-II (biotinylated, Vector laboratories) to determine the distribution of α2,3 and α2,6 linked SIAs. To detect α2,6 linked SIAs, we employed a human H1 HA derived from A/Puerto Rico/8/34 (H1PR8) (35).

We used FFPE tissues of the mammary glands of three cows and applied our library of plant-IAV- and IDV-derived glycan-binding proteins. A protein that binds a glycan is defined as a lectin, and an overview of the specificities of all lectins used is provided in the supplementary section (Fig S1). No signal was observed when we applied our antibody mix as a negative control. The classical H5 proteins, H5IN & H5VN, showed variable binding in the mammary gland, with H5IN binding significantly to cows 1 and 2 but not 3, exemplifying biological variability (Fig 2). The H5VN Y161A mutant did not bind the alveoli but bound in the mammary gland to connective tissues and blood vessels. The staining of this lectin was not as intense as expected, as cows are known for displaying 5-N-glycolyl-modified SIAs in their respiratory tract (23). The IDV HEF, on the other hand, is bound throughout the mammary gland with high intensity, confirming the abundance of 9-O-acetylated SIA.

**Figure 2.**
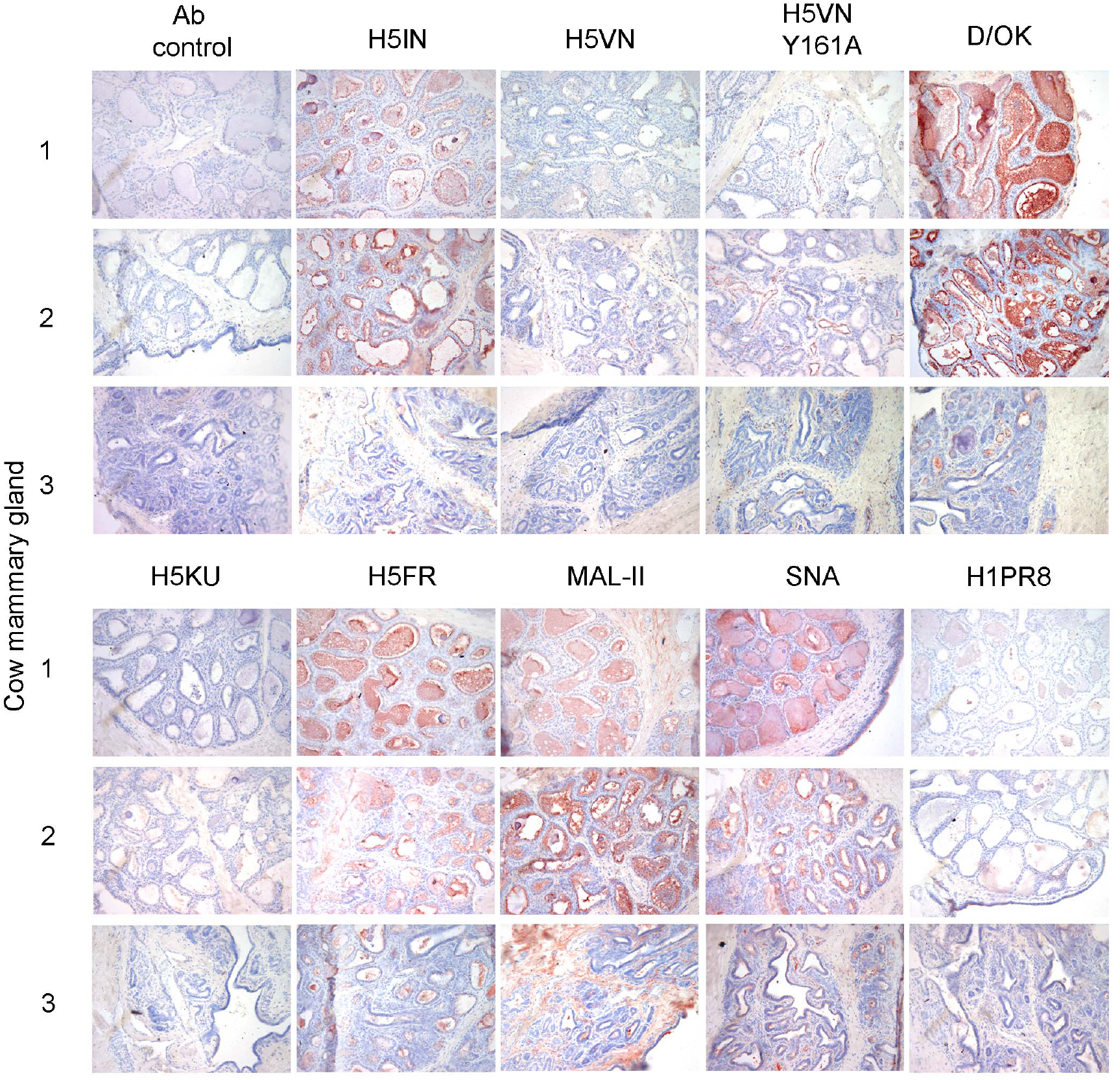
Immunohistochemical analyzes of three cow mammary glands, stained with small library of viral and plant lectins. The binding to mammary glands of three different cows was investigated for different IAV H5 proteins and a human H1 protein, as well as an IDV HEF and plant lectins MAL-II and SNA. AEC staining was used to visualize tissue binding.

**Figure 3.**
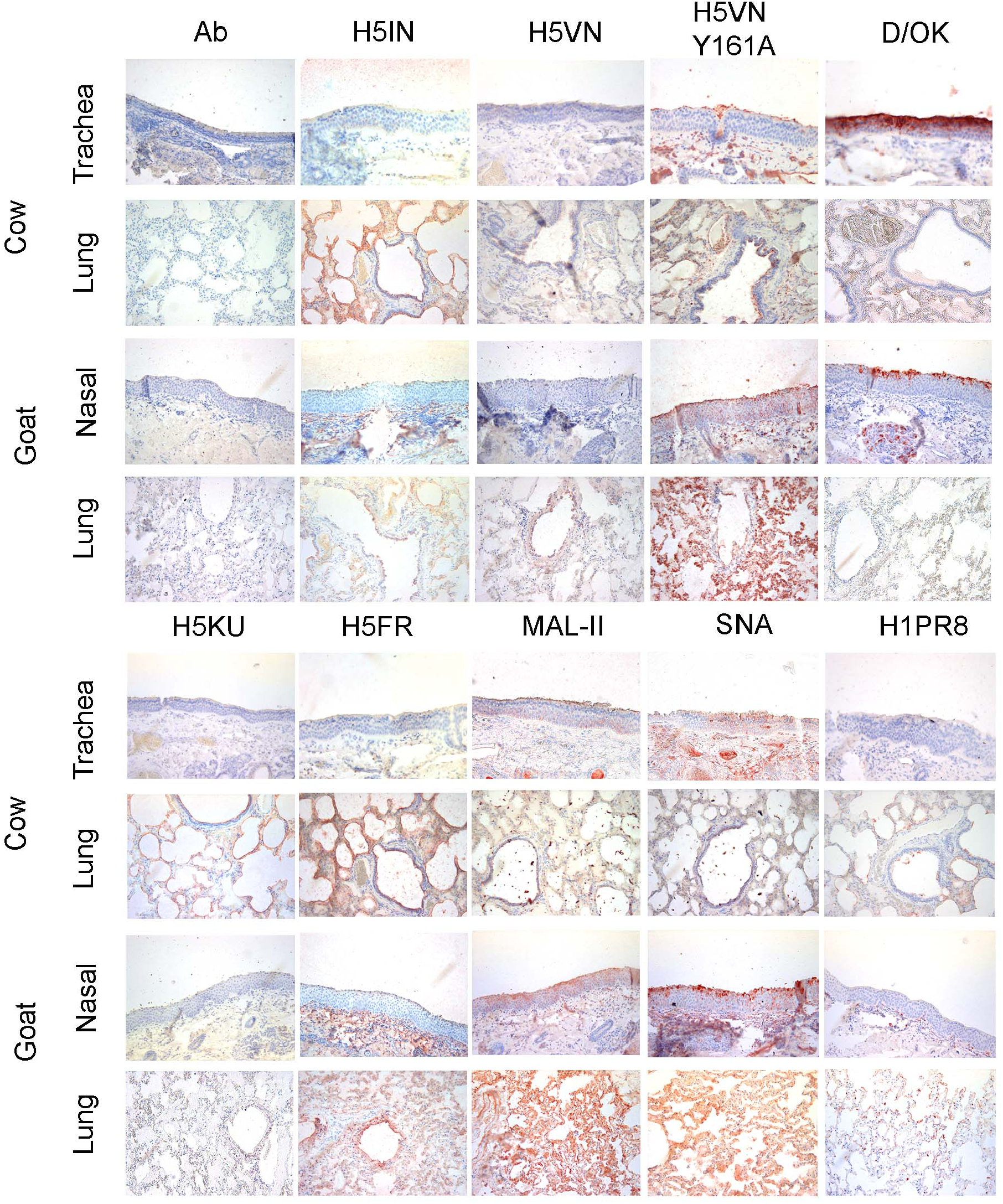
Immunohistochemical analyzes of cow and goat upper and lower respiratory tract tissues. The binding to upper (nasal or tracheal) and lower (lung) respiratory tract tissues of a cow and goat was investigated for different IAV H5 proteins and a human H1 protein, as well as an IDV HEF and plant lectins MAL-II and SNA. AEC staining was used to visualize tissue binding.

We then applied a 2.3.4.4a (H5KU) and a 2.3.4.4b HA (H5FR) and observed that the H5KU did not engage SIAs in the mammary gland, whereas the H5FR did. Intense staining is observed within the mammary glands of all three cows. A similar observation is made for both plant lectins, MAL-II and SNA. Notably, the H1 protein of a human H1N1 virus hardly bound in the mammary gland, indicating that SNA might bind α2,6 linked 5-N-glycolyl SIA (24). We confirmed the biological activity of all the lectins used using a traditional hemagglutination assay in which the proteins that bind erythrocytes form a mesh. All proteins can hemagglutinate chicken erythrocytes (Fig S2), except for the H5VN Y161A mutant, as there is no 5-N-glycolyl in avian species. H5KU also binds less avidly than the other H5 proteins but is biologically active.

Conclusively, the mammary gland of cows displays receptors for 2.3.4.4b H5 proteins and further confirms the possibility of virus binding as the first step of active replication in and transmission from this organ.

### The respiratory tract of cows and goats hardly display receptors for IAVs

In its wild waterfowl reservoir, IAVs are transmitted by the oral-fecal route, whereas in mammalian hosts, they are transmitted by the respiratory tract. Although all preliminary data points to a non-traditional transmission by milking practices, we did not want to exclude the respiratory tract of cows and goats as a possible transmission route.

We used FFPE upper (nasal or trachea) and lower (lung) respiratory tract tissues from cows and goats (N=2 and 1, respectively) and applied our library of lectins. The only H5 protein significantly binding to the upper respiratory tract of both cow and goat was the H5VN Y161A mutant, which binds to 5-N-glycolyl but that is not a receptor for circulating 2.3.4.4b H5N1 viruses. The lung tissues were bound by almost all HA proteins, including the human H1PR8, with various intensities. Furthermore, the upper respiratory tract of both cow and goat also abundantly displays 9-O-acetylated SIAs, as indicated by the high signal intensity of D/OK. Both plant lectins bound to all four tissues tested, contrasting the avian and human HAs in the tracheal and nasal tissues. This illustrates that plant-derived lectins, such as SNA and MAL-II, are poor predictors of IAV receptor distribution (36, 37).

The inability of our H5 proteins to engage in the upper respiratory tract of cows and goats and the high abundance of IAV non-receptors such as 5-N-glycolyl and 9-O-acetyl most likely exclude this transmission route. Thus, receptors are available in the lungs, and lower respiratory tract infections are often not efficiently transmitted and cause severe disease (38).

## Discussion

Here, we demonstrate that the mammary gland of cows abundantly displays avian-type receptors for circulating H5 viruses. We also show that this organ lacks human-type receptors, which contradicts a previous study that only relied on plant lectins (16). Thus, we deem the adaptation of 2.3.4.4b H5N1 viruses to human-type receptor specificity during replication in the cow mammary gland unlikely. Using direct immunohistochemical binding of H5 proteins, we demonstrate that the upper respiratory tract of goats and cows is devoid of receptors for IAV. Thus, although the zoonotic transmission of 2.3.4.4b H5N1 viruses to ruminants is unprecedented, we would suggest it does not pose an immediate pandemic risk, as there is no need to adapt to human-type receptors. However, the continuous and widespread circulation of these zoonotic viruses in primary livestock is a concern. Importantly, in human infections, the virus is almost only isolated from eye swabs (39). Although conjunctivitis in humans caused by AIV has been observed before and can be modeled in the ferret model (40, 41), after such ocular inoculation, the virus could further spread to the upper respiratory tract, in which H5N1 viruses might adapt to human-type receptors.

We focused our attention on the display of sialylated receptors for IAV HA proteins. We have previously shown that the receptor specificity of HA proteins is very similar to that of whole viruses, although with a lower degree of multivalency, leading to lower signals (24). However, we can easily observe binding in the cow’s mammary gland. Using multiple cows was vital as the signals in cow 3 are significantly lower, which probably presents natural variation. We would like to emphasize our use of an IDV HEF protein that stains almost all tested tissues with high intensity. Cows are the reservoir for IDVs, and the abundant display of 9-O-acetylated SIAs could be a block binding of IAV. Finally, we would like to point out that sole use of plant lectins to study the differential display of avian- and human-type receptors (36, 37), leads to an over-interpretation as their binding profile is more promiscuous.

We expect that our studies into the display of SIA modifications in ruminants will aid in our understanding of how different IAV viruses with distinct glycan specificities infect and transmit. Indeed, a recent unofficial report paints a picture of severe infections in cows, which might be related to lower respiratory tract infections, as ample receptors are available there (42). The glycome of horses and pigs are not expected to differ greatly from cows, as both abundantly display 5-N-glycolyl and 9-O-acetyls. While horses are hardly ever infected with avian IAVs, pigs are, and experimental infection of pigs with 2.3.4.4b viruses have shown that they are susceptible (43, 44), so a question is why we do not observe natural 2.3.4.4b H5N1 infections in pigs.

Finally, MS-based analyses are essential to further map the sialome, and controlled infection studies are desperately needed to understand this zoonotic event.

## Acknowledgments

This research was made possible by funding from ICRAD, an ERA-NET co-funded under the European Union’s Horizon 2020 research and innovation programme (https://ec.europa.eu/programmes/horizon2020/en), under Grant Agreement n°862605 (Flu-Switch) to, and a Mizutani Foundation for Glycoscience Research Grant 2023 to Robert P. de Vries.

We thank the staff of the Veterinary Pathology Diagnostic Centre of Utrecht University for their assistance in collecting the tissues.

## Materials and methods

### Expression and purification of trimeric influenza A hemagglutinins

Recombinant trimeric IAV hemagglutinin ectodomain proteins (HA) were cloned into the pCD5 expression vector (an example is addgene plasmid #182546) in frame with a GCN4 trimerization motif (KQIEDKIEEIESKQKKIENEIARIKK), a super folder GFP or mOrange2 (45) and the Twin-Strep-tag (WSHPQFEKGGGSGGGSWSHPQFEK); IBA, Germany). The trimeric HAs were expressed in HEK293S GnTI(-) cells with polyethyleneimine I (PEI) in a 1:8 ratio (μg DNA:μg PEI) for the HAs as previously described. The transfection mix was replaced after 6 hours by 293 SFM II suspension medium (Invitrogen, 11686029), supplemented with sodium bicarbonate (3.7 g/L), primatone RL-UF (3.0 g/L, Kerry, NY, USA), glucose (2.0 g/L), glutaMAX (1%, Gibco), valproic acid (0.4 g/L) and DMSO (1.5%). According to the manufacturer’s instructions, culture supernatants were harvested five days post-transfection and purified with sepharose strep-tactin beads (IBA Life Sciences, Germany).

### Protein histochemical tissue staining

Sections of formalin-fixed, paraffin-embedded cow (*Bos taurus*) and goat (*Capra aegagrus hircus*) were obtained from the Division of Pathology, Department of Biomolecular Health Sciences, Faculty of Veterinary Medicine of Utrecht University, the Netherlands. In the figures, representative images of at least two individual experiments are shown. Protein histochemistry was performed as previously described (46, 47). In short, tissue sections of 5μm were deparaffinized and rehydrated, after which antigens were retrieved by heating the slides in 10 mM sodium citrate (pH 6.0) for 10 min. Endogenous peroxidase was inactivated using 1% hydrogen peroxide in MeOH for 30 min at RT. Tissues were blocked at 4°C using 3% BSA (w/v) in PBS for at least 90 minutes. Subsequently, slides were stained for 90 minutes with 10μg/mL solution of precomplexed proteins of interest. HAs were precomplexed with human α-strep-tag primary antibody and goat-α-mouse-HRP secondary antibody at 4:2:1 molar ratio as previously described (12). For biotinylated plant lectins, we used streptavidin-HRP at 4:1 molar ratio. We used 3-amino-9-ethylcarbazole (AEC) (Sigma-Aldrich, Steinheim, Germany) to visualize protein binding. Tissue sections were then counterstained with hematoxylin and mounted with coverslips using AquaTex (Merck).

---

**Figure S1.**
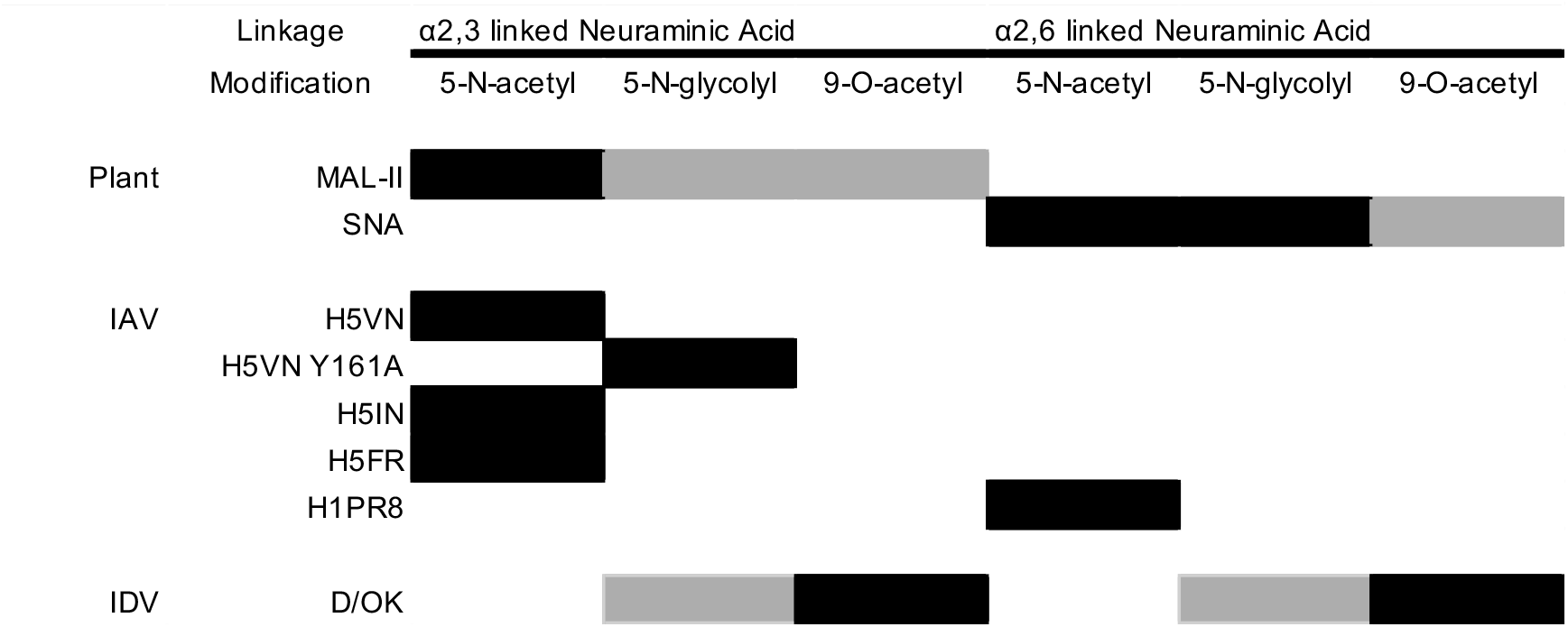
Overview of lectin specificity to differentially linked un- and modified SIA. Black boxes indicate core specificities, gray boxes indicate possible ligands.

**Figure S2.**
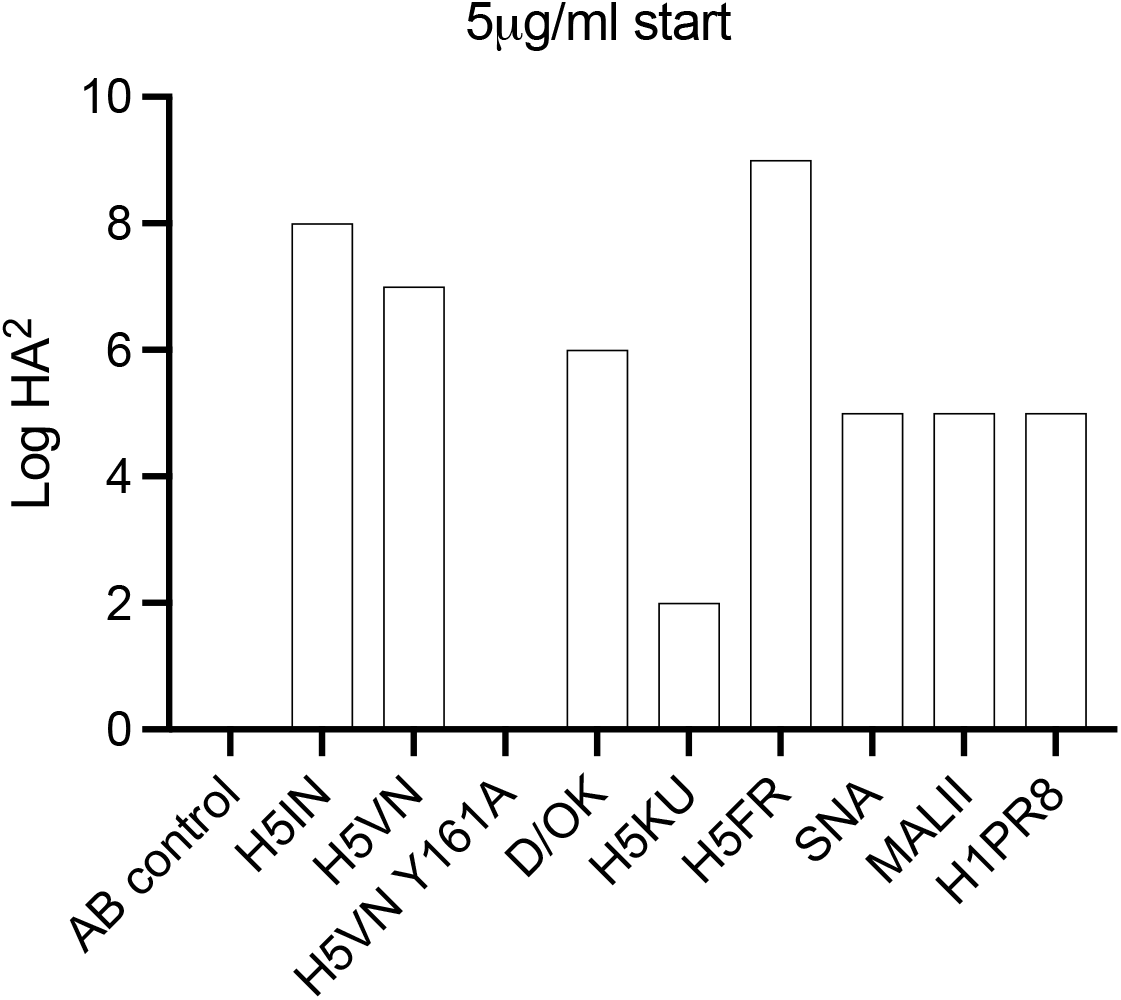
Hemagglutination assay with chicken erythrocytes with the lectins used in this study.

## Notes

### Competing Interest Statement

The authors have declared no competing interest.

### Summary of Updates

We needed to update the abstract, but I forgot to change it in submission program as well. Both PDF and abstract online are good now. Many thanks and sorry for the inconvenience.

